## Supplemental Figure for "Scalable primordial germ cell-like-cell platform for functional genomics identifies epigenetic fertility modifiers"

### Scalable Generation of Primordial Germ Cell-like Cells for Functional Perturbation Screens Identifies Key Epigenetic Genes

Liangdao Li[1#], Jingyi Gao[1], Dain Yi[1], Alex P. Sheft[1], John C. Schimenti[1,2*] and Xinbao Ding[1#]

1 Cornell University, College of Veterinary Medicine, Department of Biomedical Sciences, Ithaca, NY 14853.

2 Cornell University, College of Agriculture and Life Sciences, Department of Molecular Biology and Genetics, Ithaca, NY, 14853, USA

[#] Contributed equally

**Supplementary Figures 1~21**

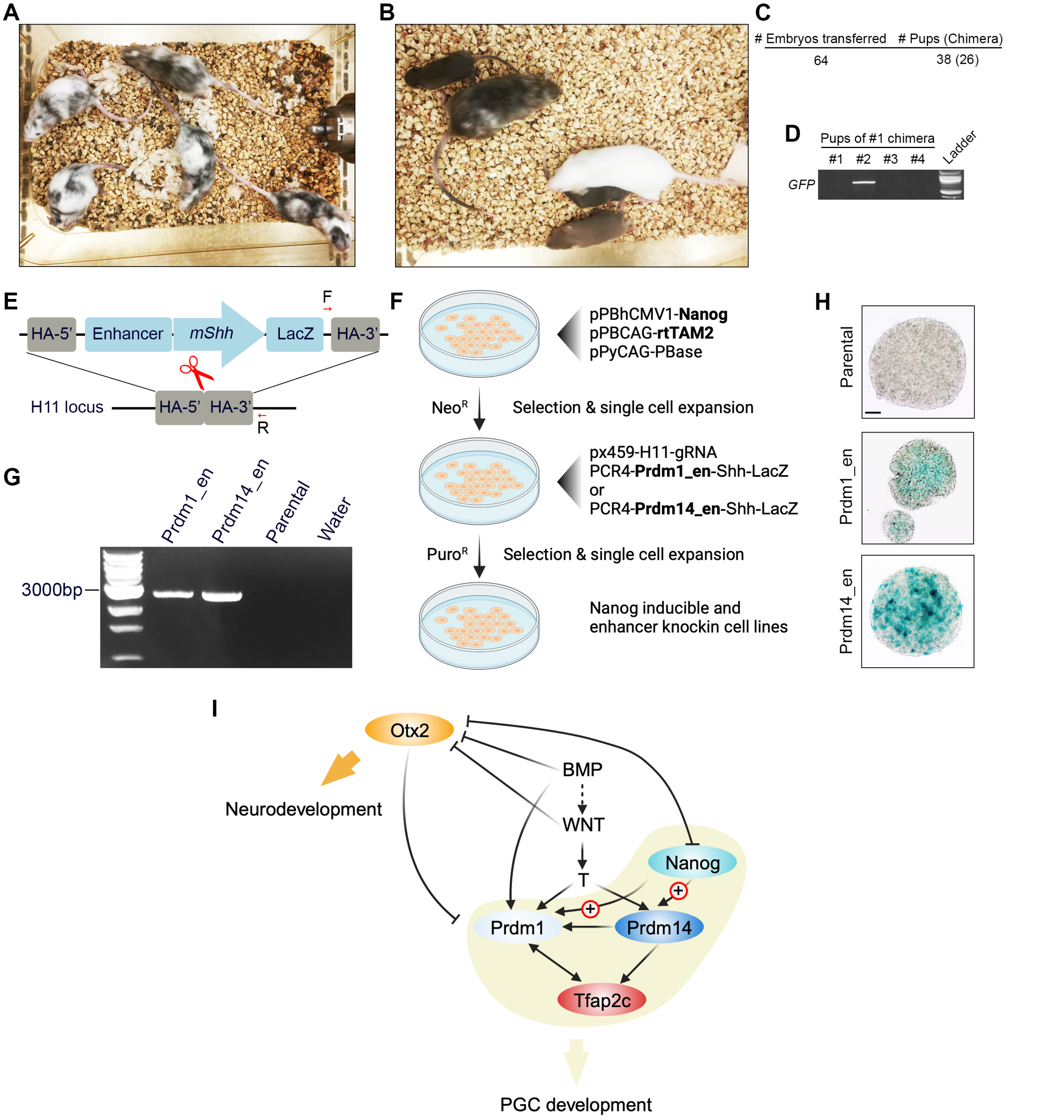


**Figure S1. Assessing the germline competence of Stella-eGFP ESCs and confirming NANOG's regulation of *Prdm1* and *Prdm14*.** A. Chimeras generated by Stella-eGFP ESCs. B. Representative of germline transmission of chimeras from Stella-eGFP ESCs. C. Summary of chimeric mice generated by blastocyst microinjection. D. Genotyping of pups to detect transgene originating from Stella-eGFP ESCs. E. Scheme for CRISPR/Cas9- assisted transgene integration into the H11 "safe harbor" locus. F. Overview of how ESCs were generated that contain NANOG-responsive reporters (*Prdm1* and *Prdm14* enhancers upstream of a *Shh* basal promoter) at the H11 locus. G. PCR verification of clones harboring knock-in vectors. H. Representative β-gal staining of EBs. I. Schema illustrating key TFs acting during PGC development. Scale bar in H represents 50 μm.

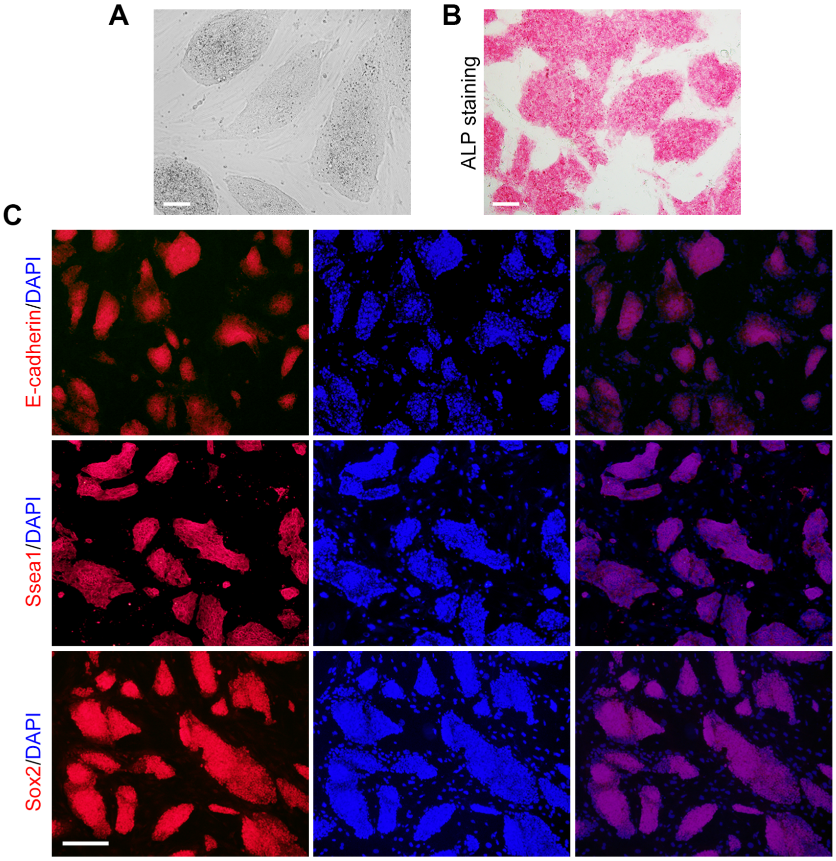


**Figure S2. Reprogramming properties of 4TF-induced PGCLCs.** A. Morphology of EGCs derived from 4TF-induced PGCLCs (Stella-eGFP$^{+}$ cells) at day 6. B. ALP (alkaline phosphatase) staining of EGCs. C. IF of E-cadherin, SSEA1 and SOX2 in EGCs. Scale bars in A and B = 50μm and in C = 100μm.

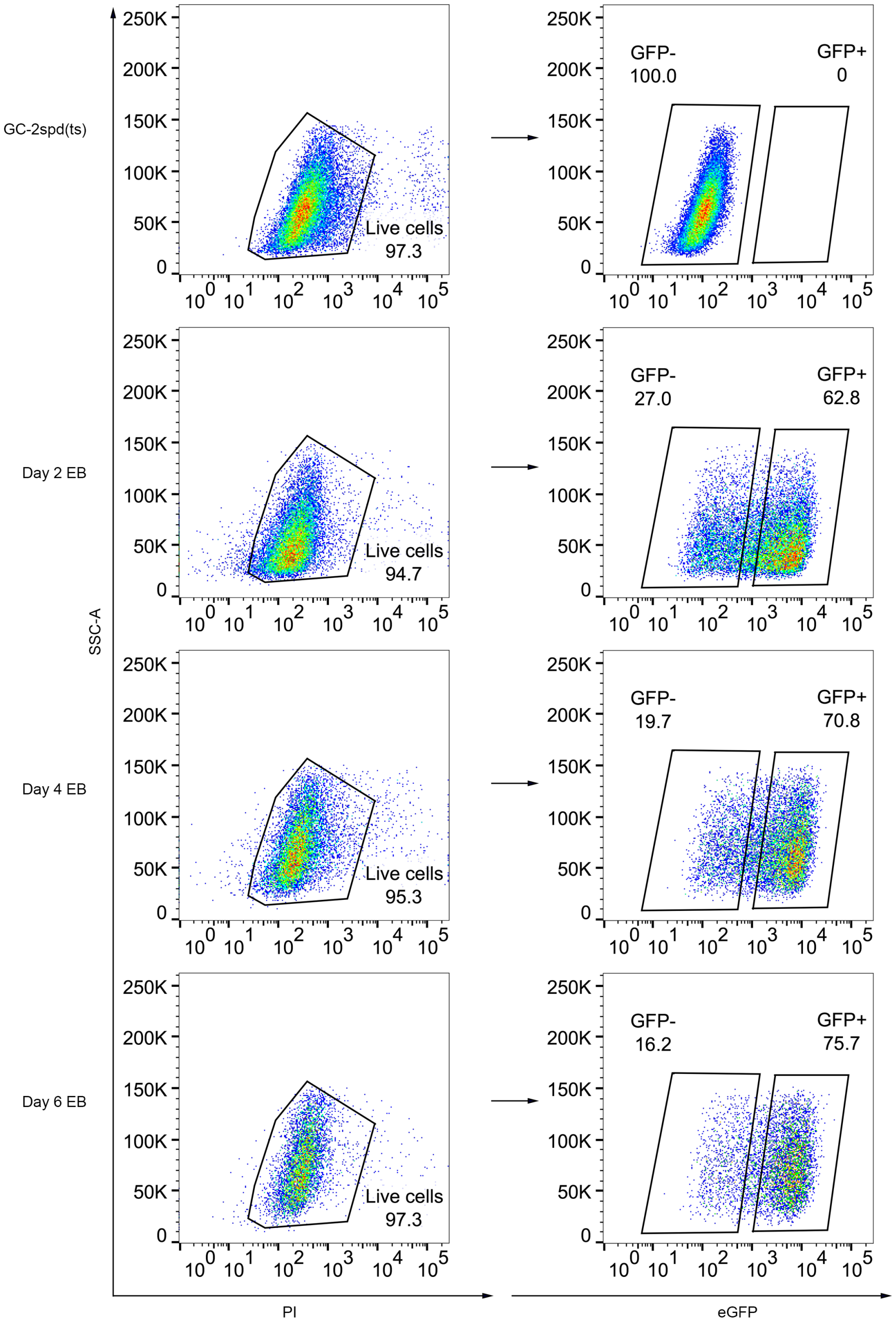


**Figure S3. FACS gating strategy for the isolation of PGCLCs (Stella-GFP+) from EBs.** GC-2spd(ts) cells served as $GFP^{-}$ reference. Gating for live cells was performed using propidium iodide (PI).

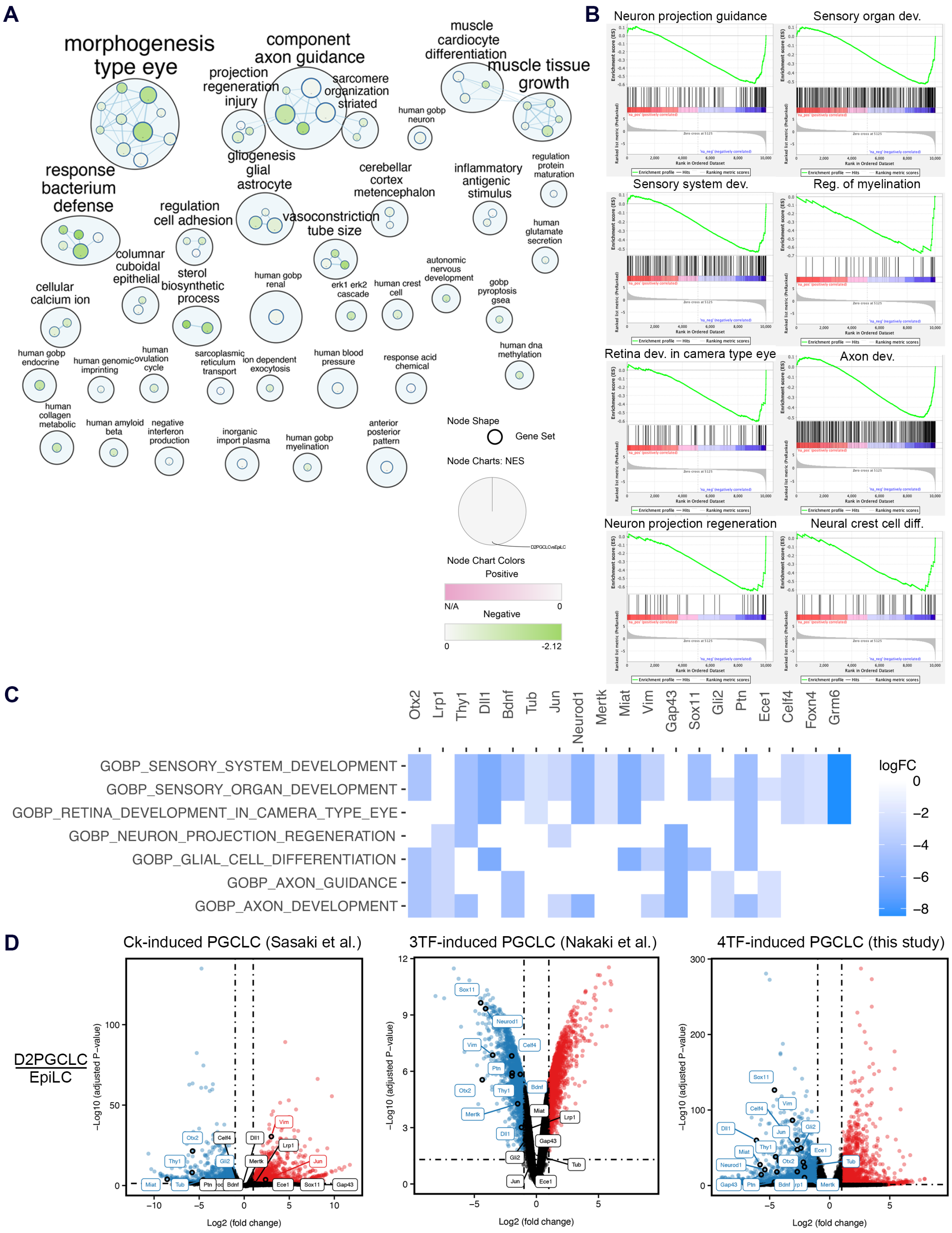

**Figure S4. Neurodevelopment pathways suppressed in 4TF-induced PGCLCs.** A. Network map illustrating enrichment of GOBP (Gene Ontology Biological Process) in EpiLCs vs. D2PGCLCs. Gene sets (nodes) are connected based on shared edges. Nodes downregulated in EpiLCs are highlighted in green. The size of each node corresponds to the percentage of enriched genes in the set, and color intensity indicates the degree of expression change. Blue lines represent shared gene overlap between sets. Clusters of gene sets with related functions are circled and labeled to emphasize prevalent biological functions. B. Suppression of neurodevelopmental processes in EpiLCs vs. D2PGCLCs. C. Heatmap of GO enrichment analysis for a subset of genes related to neurodevelopment. Displayed are log2 fold change (logFC) values between D2PGCLCs and EpiLCs. White indicates no association with the corresponding pathway. D. Volcano plot comparing transcriptomes of Ck-, 3TF- and 4TF- induced PGCLCs to EpiLCs. Dashed lines represent significance thresholds. Selected genes involved in neurodevelopment are highlighted.

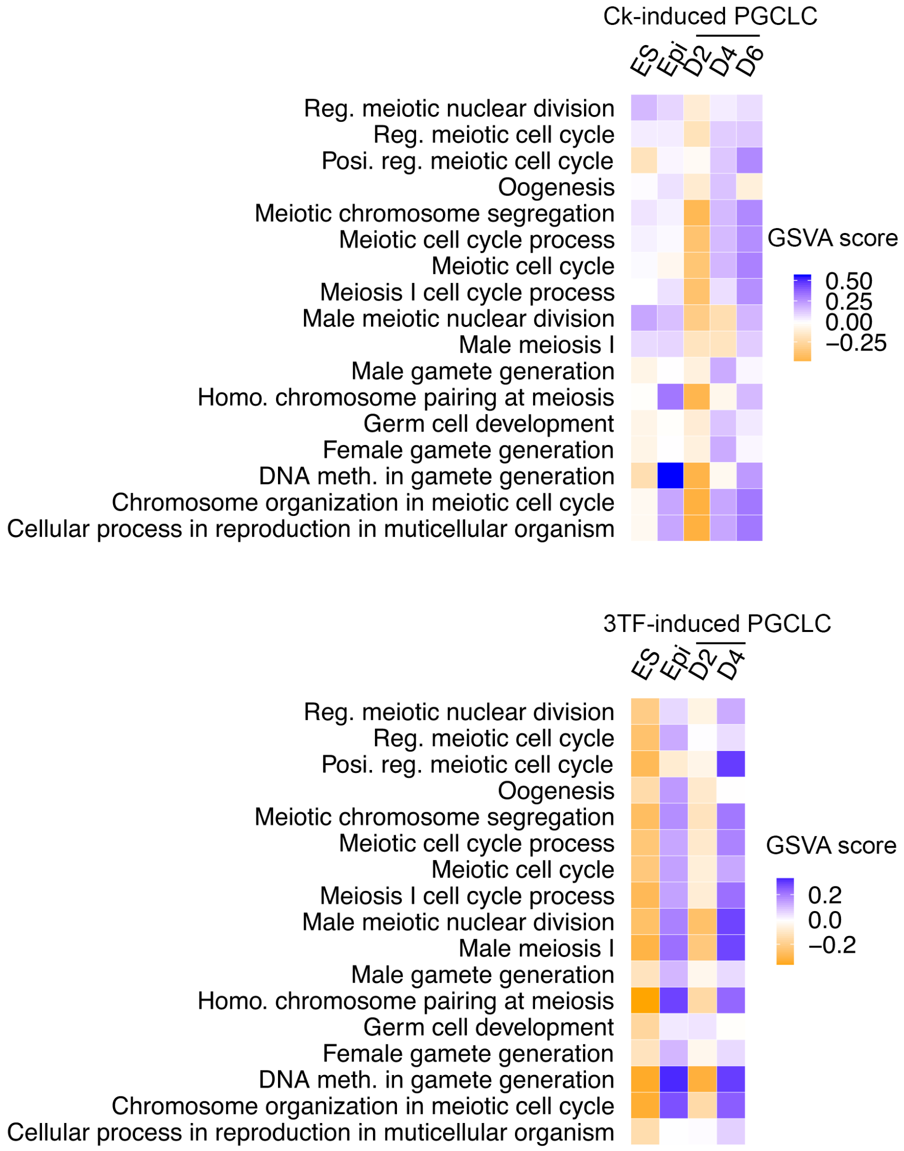


**Figure S5. Heatmaps showing the average GSVA enrichment score of selected germ cell development pathways in Ck- and 3TF- induced PGCLCs**

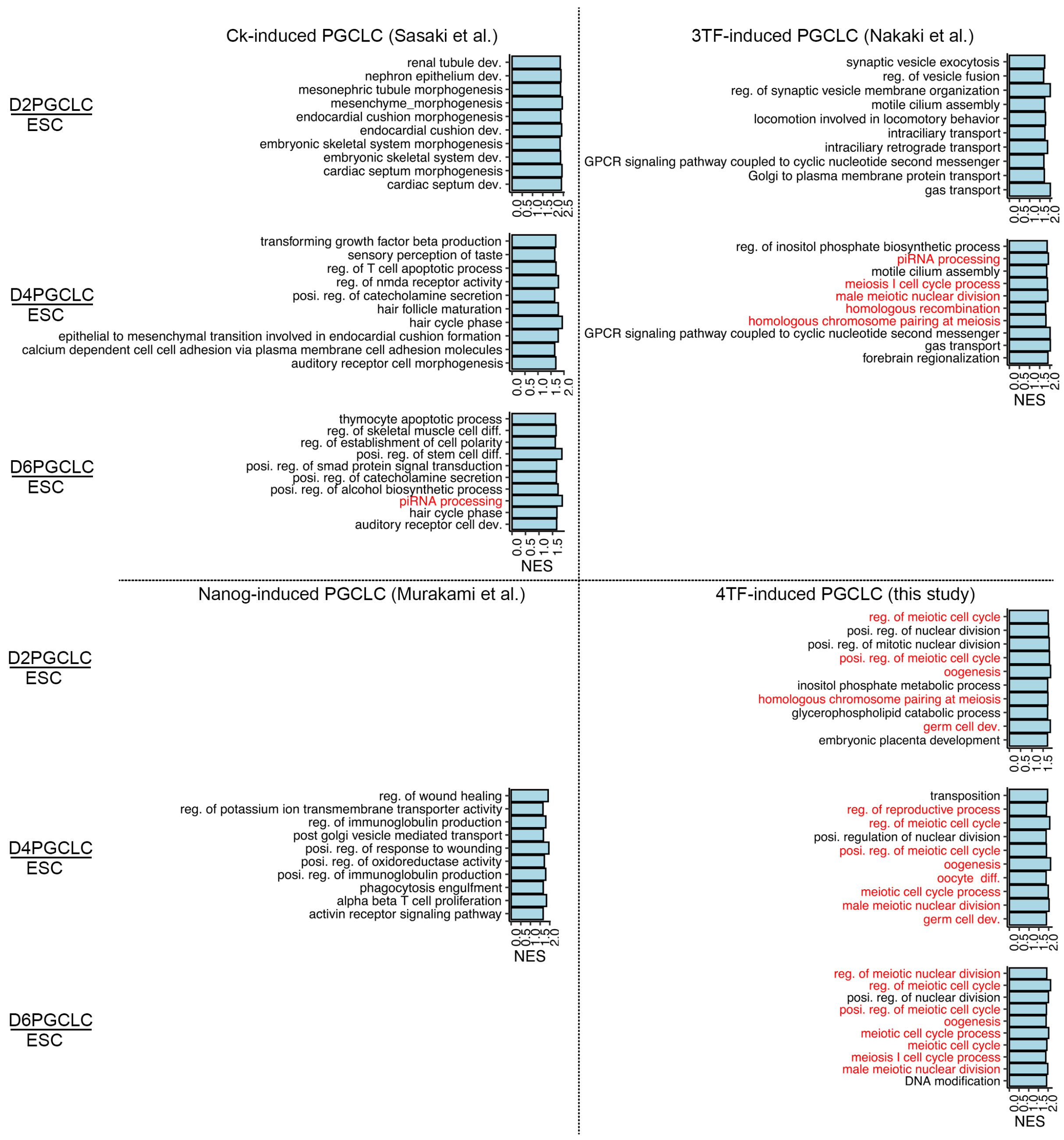


**Figure S6. Top 10 enriched biological process GO terms in Ck-, 3TF-, Nanog- and 4TF-induced PGCLCs identified by GSEA.** Terms of germ cell development (highlighted by red) was highly enriched in 4TF-induced PGCLCs. All the p-value of these GO terms were < 0.05. NES, normalized enrichment score.

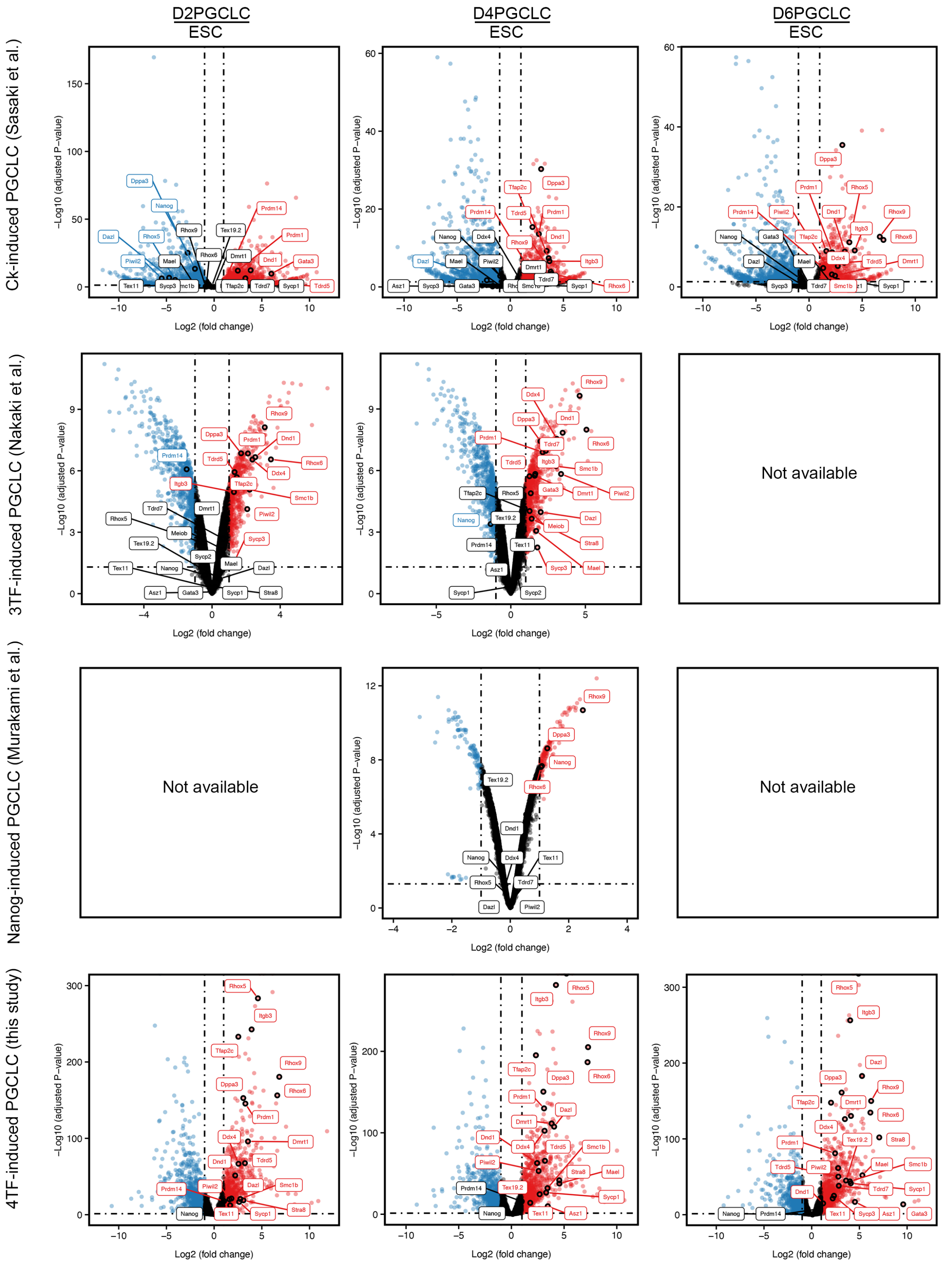


**Figure S7. Volcano plot comparing transcriptomes of Ck-, 3TF-, Nanog- and 4TF-induced PGCLCs to ESC.** Dashed lines represent significance thresholds. Selected genes involved in germ cell development are highlighted.

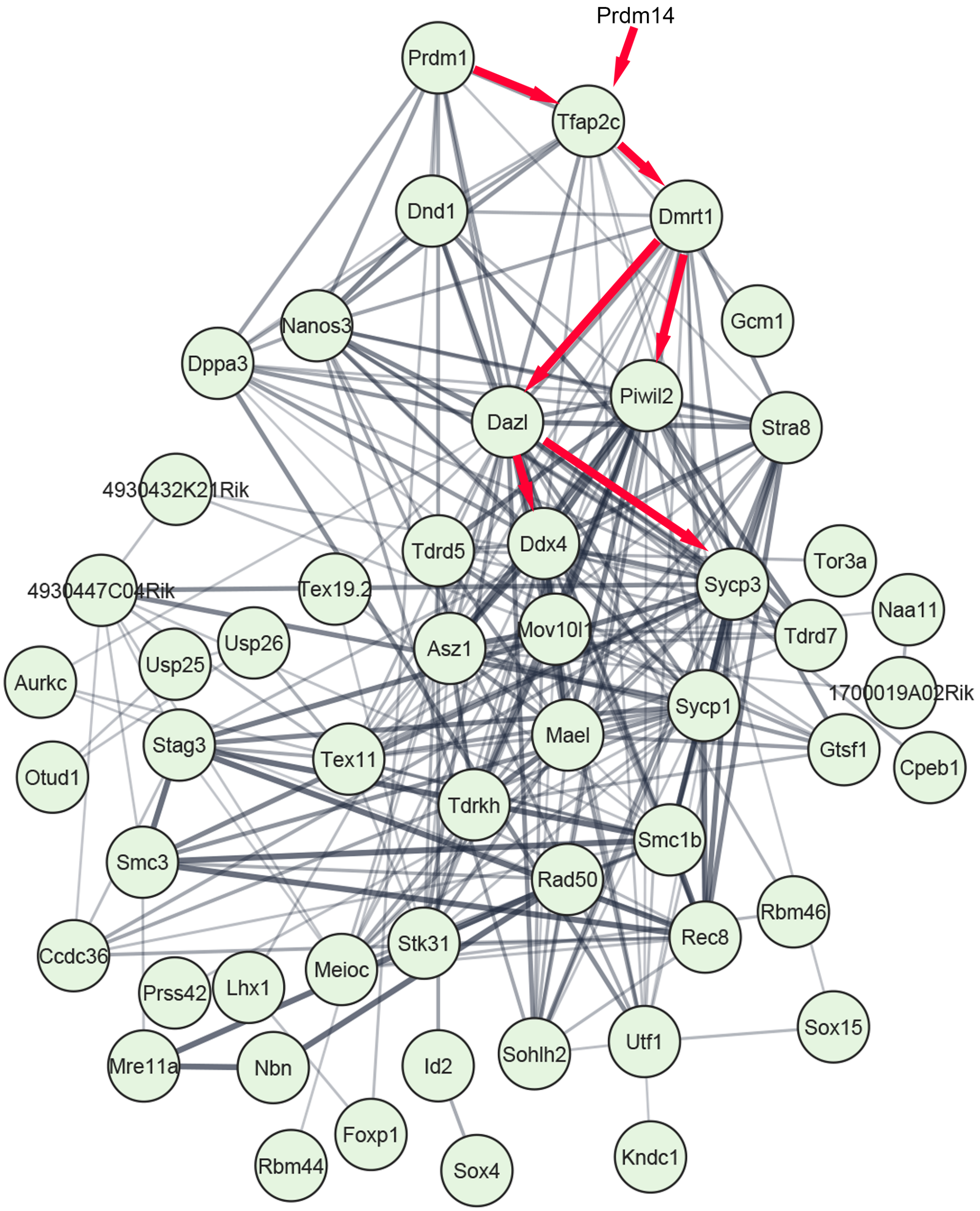


**Figure S8. STRING analysis of upregulated DEGs in PGCLCs compared to ESCs.** *Prdm14* was omitted from the network construction because the log2(FC) of PGCLCs vs ESCs was <2. Red arrows indicate the transition from early to late PGC.

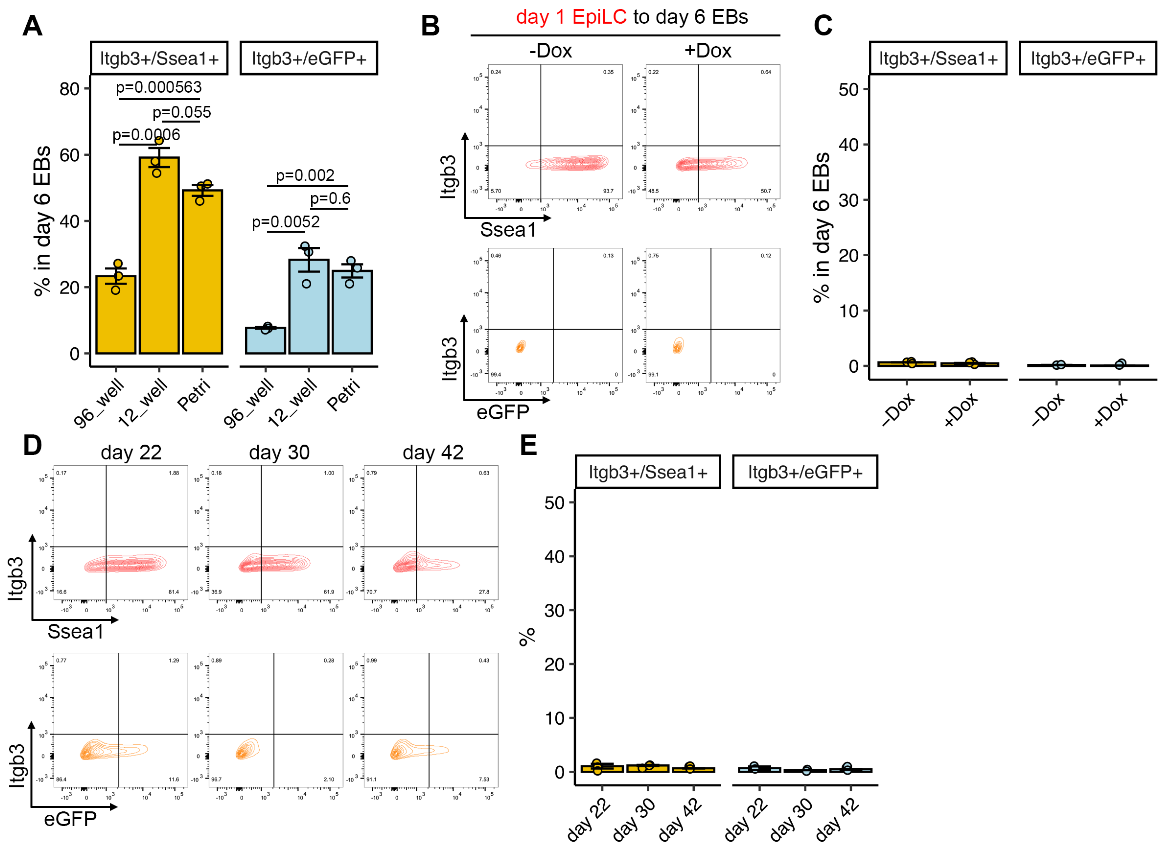


**Figure S9. Differentiation of Nanog-induced PGCLCs.** A. Quantification of PGCLC populations induced by Nanog in day 6 EBs at different plates/ dish. B. Representative FACS pattern of Itgb3⁺/Ssea1⁺ and Itgb3⁺/eGFP⁺ cells after inducing *Nanog* alone or not in day 1 EpiLCs for 6 days' suspension culture. C. Quantification of PGCLC populations in day 6 EBs with or without Dox treatment. D. Representative FACS pattern of Itgb3⁺/Ssea1⁺ and Itgb3⁺/eGFP⁺ cells after long-term culture in *Nanog*-inducible system. C. Quantification of PGCLC populations at different time points of long-term cultured EBs. Data in A, C and E are represented as the mean ± SEM. Data in A were analyzed using one-way ANOVA with Tukey's *post hoc* test.

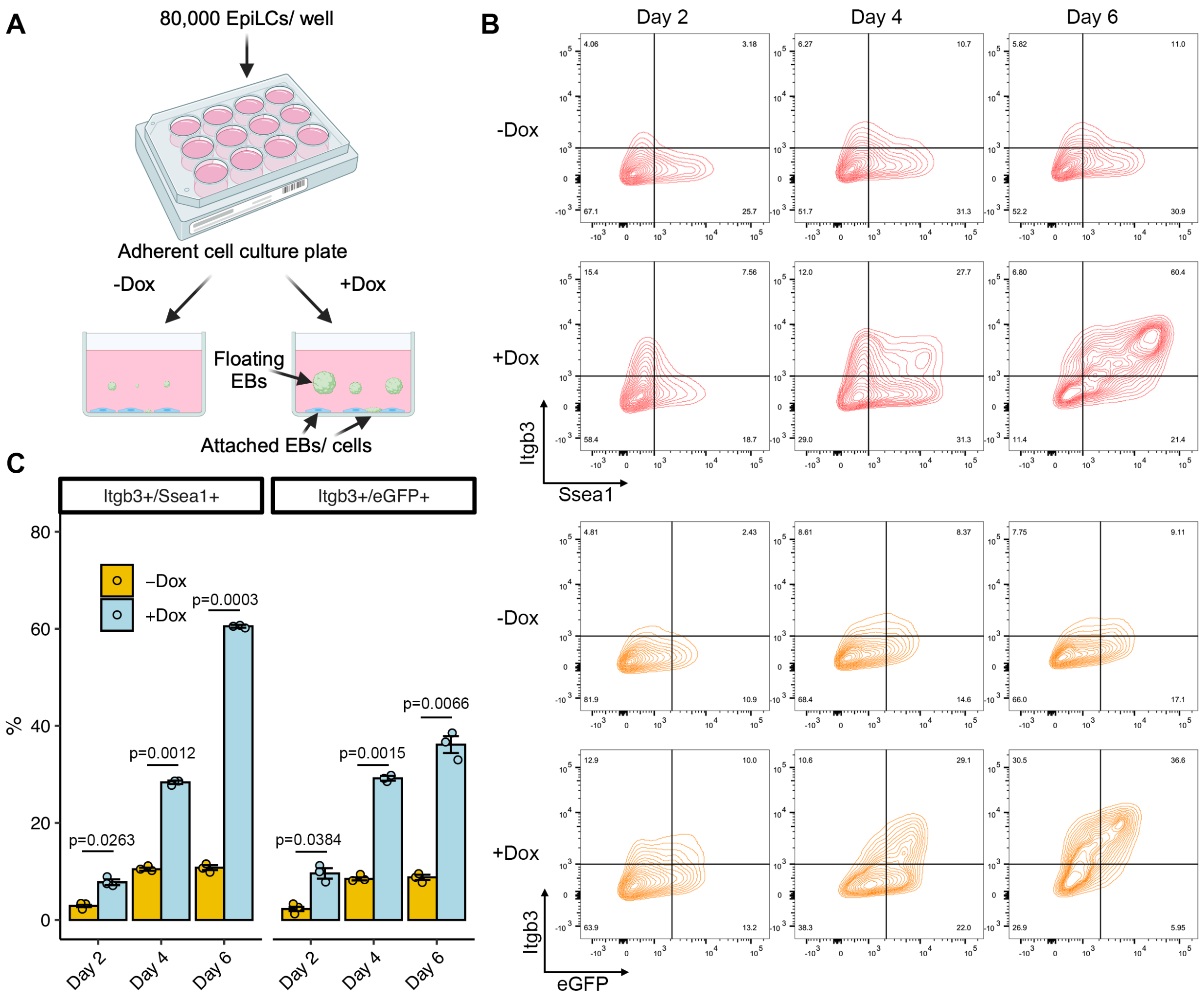


**Figure S10. Differentiation of 4TF-induced PGCLCs in adherent cell culture plates.** A. Schema of PGCLC differentiation; both floating EBs and attached cells were observed. B. Representative FACS pattern of $Itgb3^{+}/Ssea1^{+}$ and $Itgb3^{+}/eGFP^{+}$ cells after EpiLCs were treated by Dox or not during the 6-day period. The floating and attached cells were grouped for analysis. C. Quantification of PGCLC populations. Data in C are represented as the mean ± SEM and analyzed using a two-tailed paired *t* test.

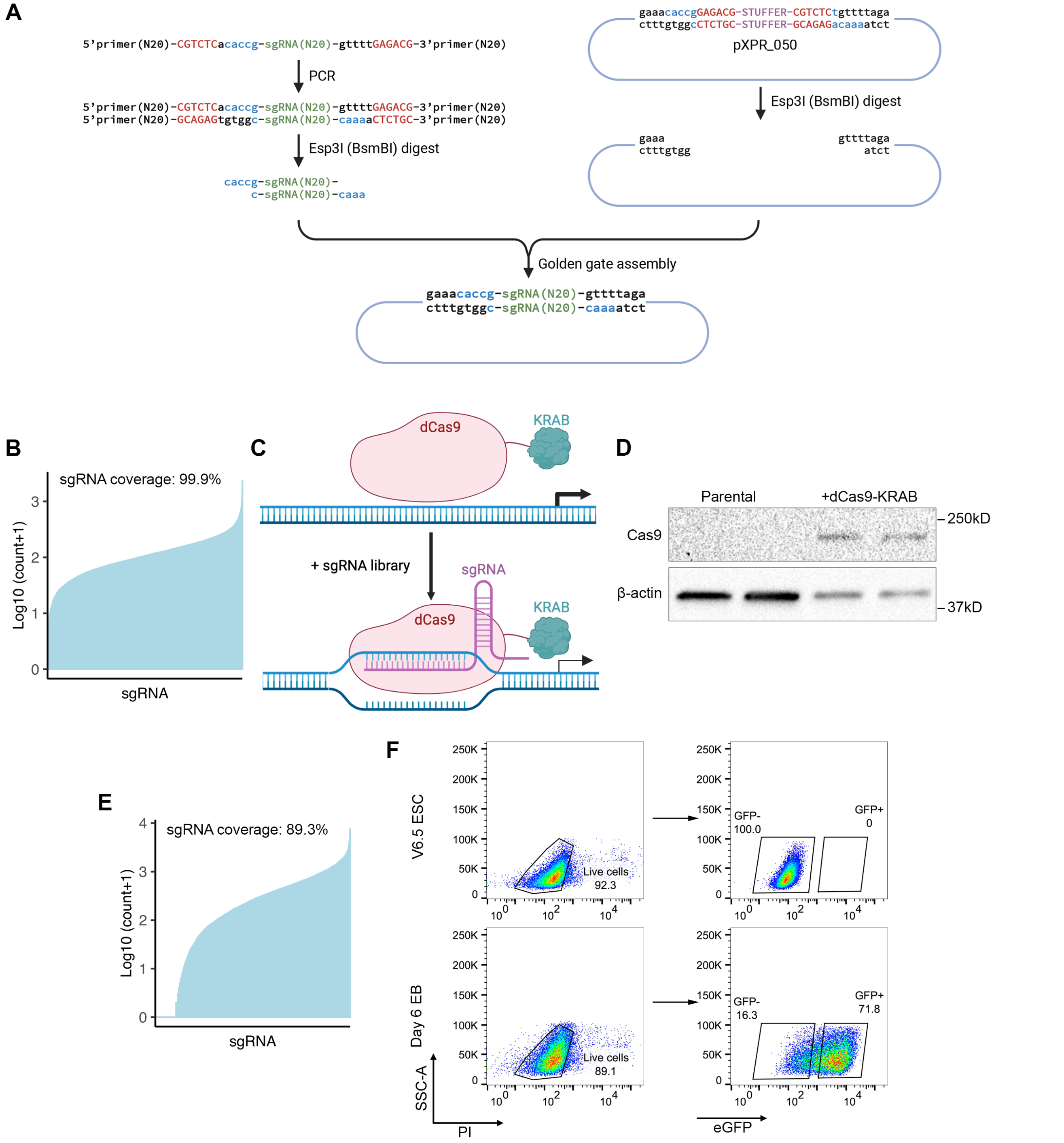


**Figure S11. Library construction.** A. Plasmid cloning of CRISPRi library. B. sgRNA coverage of the synthesized CRISPRi plasmid pool. C. Schematic illustration of CRISPRi using dCas9 and a KRAB repression domain programmed by sgRNA. D. dCas9 and beta actin protein levels of parental ESCs and ESCs stably transduced with dCas9-KRAB. E. Coverage of sgRNA vector integrations after viral infection into ESCs. F. FACS of Stella-eGFP$^{+}$ and Stella-eGFP$^{-}$ cells from day 6 dispersed EBs. PI, propidium iodide. V6.5 ESC, non-transgenic ESC line.

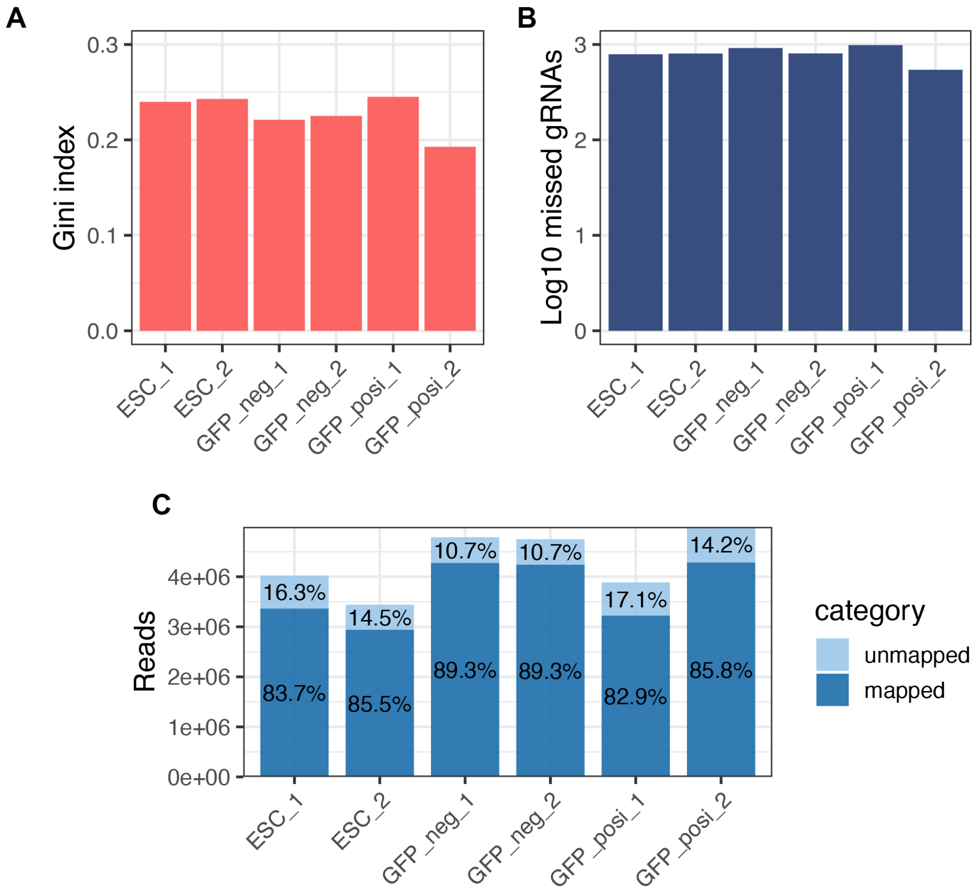


**Figure S12. CRISPRi library characterization.** (A) The Gini index measures the evenness of sgRNA read count. (B) Normalized number of missed sgRNA read count. (C) Number of reads and percentage of unmapped/mapped reads. All samples are from the CRISPRi screen dataset generated from ESCs, Stella-eGFP$^{-}$ (GFP_neg) and Stella-eGFP$^{+}$ (GFP_posi) cells. Data were generated using Screen Processing Tools pipeline.

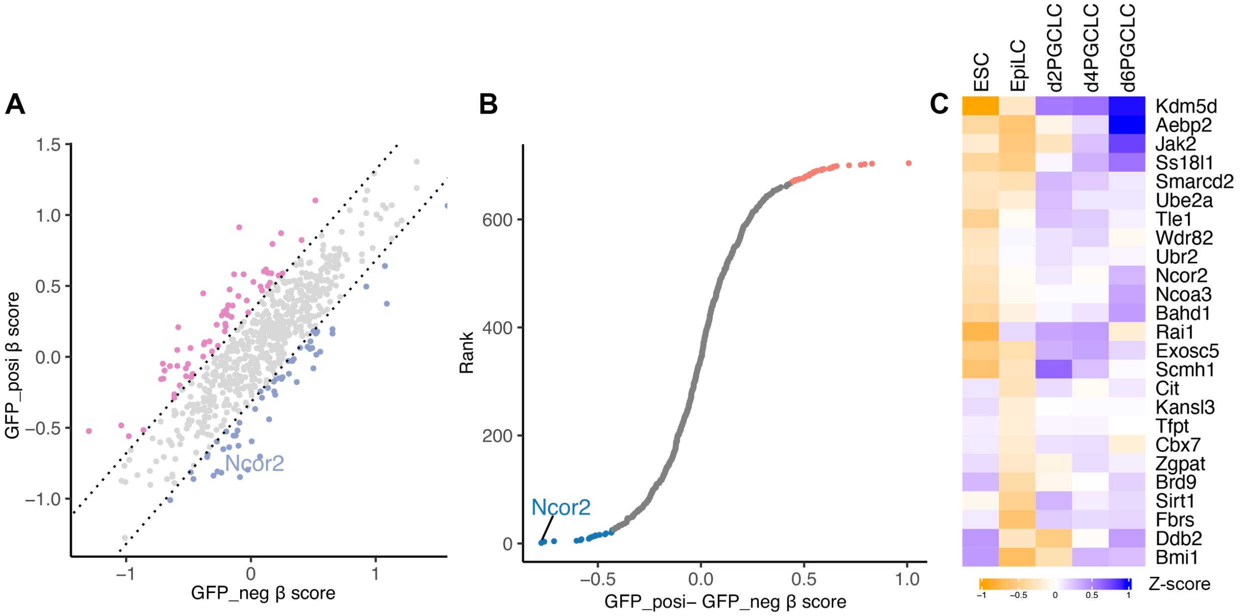


**Figure S13. CRISPRi screen epigenetic genes affecting PGCLC differentiation.** A. Scatterplot of Stella-eGFP$^{+}$ and Stella-eGFP$^{-}$ β scores of each gene. The β scores were normalized using non-targeting sgRNA sequences. The two dashed lines indicates ± 1 S.D. of the differences between the Stella-eGFP$^{+}$ and Stella-eGFP$^{-}$ cells β scores. B. Rank plot of the differential β scores, calculated by subtracting Stella-eGFP$^{-}$ β scores from Stella-eGFP$^{+}$ β scores. The color scheme of the dots is the same as in A. C. Gene expression heatmap of 25 candidate genes with β scores and exhibited increased expression during PGCLC differentiation compared to ESC and EpiLC stages.

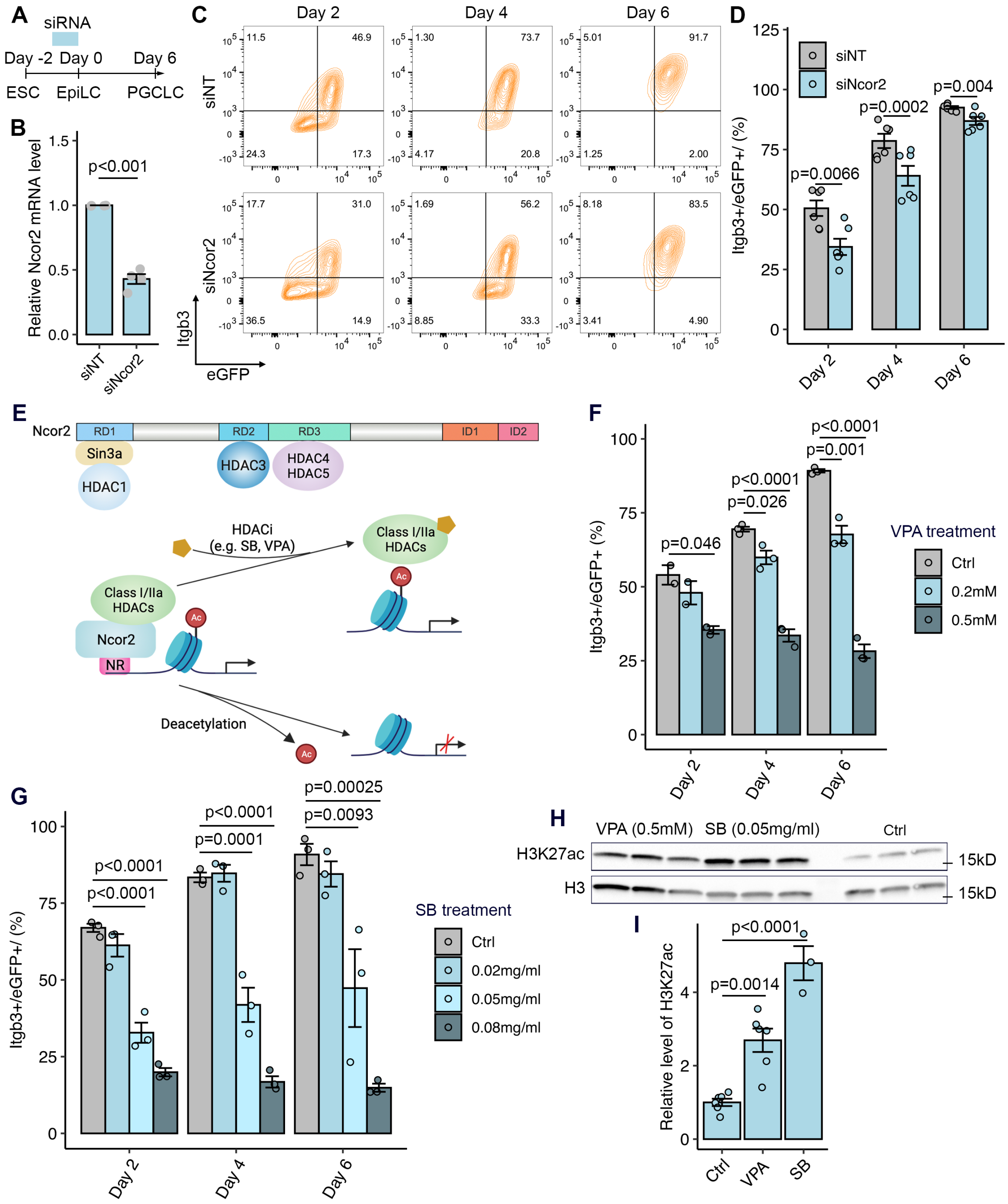


**Figure S14. NCOR2 deficiency and HDAC inhibitors suppress PGCLC differentiation *in vitro.*** A. Experimental schema showing siRNA treatment. B. Effectiveness of *Ncor2* siRNA on day 2 after PGCLC induction. mRNA levels quantified by qRT-PCR. siNT, non-targeting siRNA. C. Relative population density in EBs treated with siNT or siNcor2. siNcor2, siRNA targeting *Ncor2*. D. Percentages of Itgb3$^{+}$/eGFP$^{+}$ cells in knockdown (siNcor2) vs control (siNT) groups at the indicated stages after transfection of small RNAs. eGFP, Stella-eGFP. E. Structure of NCOR2 and interaction with HDACs (upper panel). Impact of HDAC inhibitors on NCOR2-HDAC complexes (lower panel). F. Percentage of Itgb3$^{+}$/eGFP$^{+}$ populations within EBs treated with water (Ctrl) or VPA as assessed by flow cytometry. G. Same as “F”, except for SB treatment. H. Western blot analysis of H3K27ac and H3 protein levels in day 2 Stella-eGFP$^{+}$ cells treated with HDACi. I. Quantification of relative H3K27ac protein levels normalized to H3. Data in B, D, F, G and I are represented as the mean ± SEM. Data in B were analyzed using a two-tailed paired *t* test, and data in D, F, G and I were analyzed using one-way ANOVA with Tukey’s *post hoc* test.

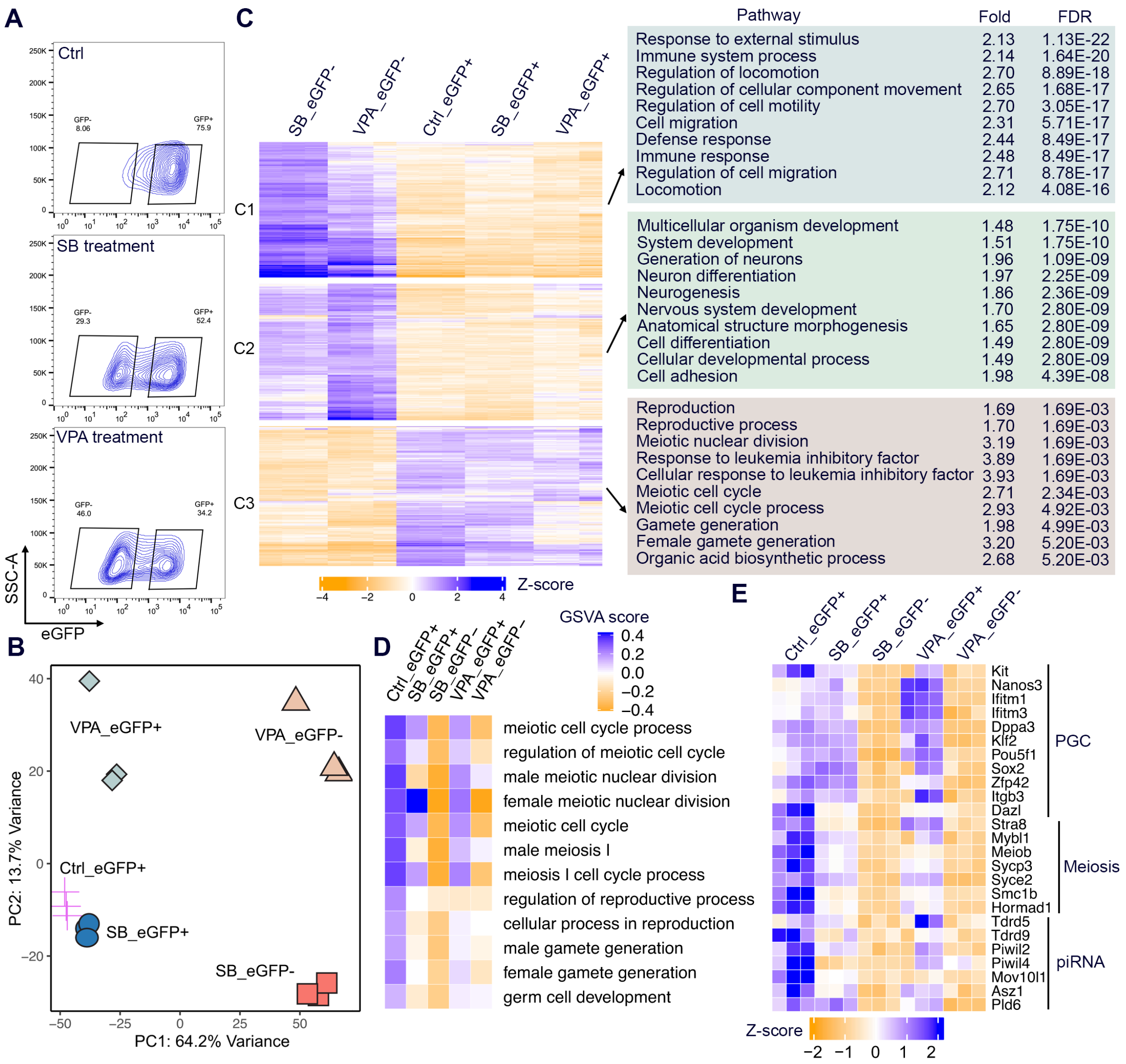


**Figure S15. Downregulation of germline networks by HDAC inhibitors.** A. Flow sorting gates for Stella-eGFP$^{+}$ and Stella-eGFP$^{-}$ cells from day 6 EBs used for RNA-seq. B. PCA of transcriptomes of sorted cells of indicated groups. SB, sodium butyrate; VPA, valproic acid; Ctrl, control untreated. C. Heatmap of k-Means clustering of variably expressed genes in SB, VPA and Ctrl cells (n = 2,000; k = 3). Genes were grouped into 3 clusters (“C1-3”) based on expression similarity. Top enriched GO terms for the genes in each cluster shown with fold enrichment (Fold) and false discovery rate (FDR). D. Heatmap showing the average GSVA enrichment score of selected germline development pathways. E. Heatmaps of normalized RNA-seq reads for selected germline genes.

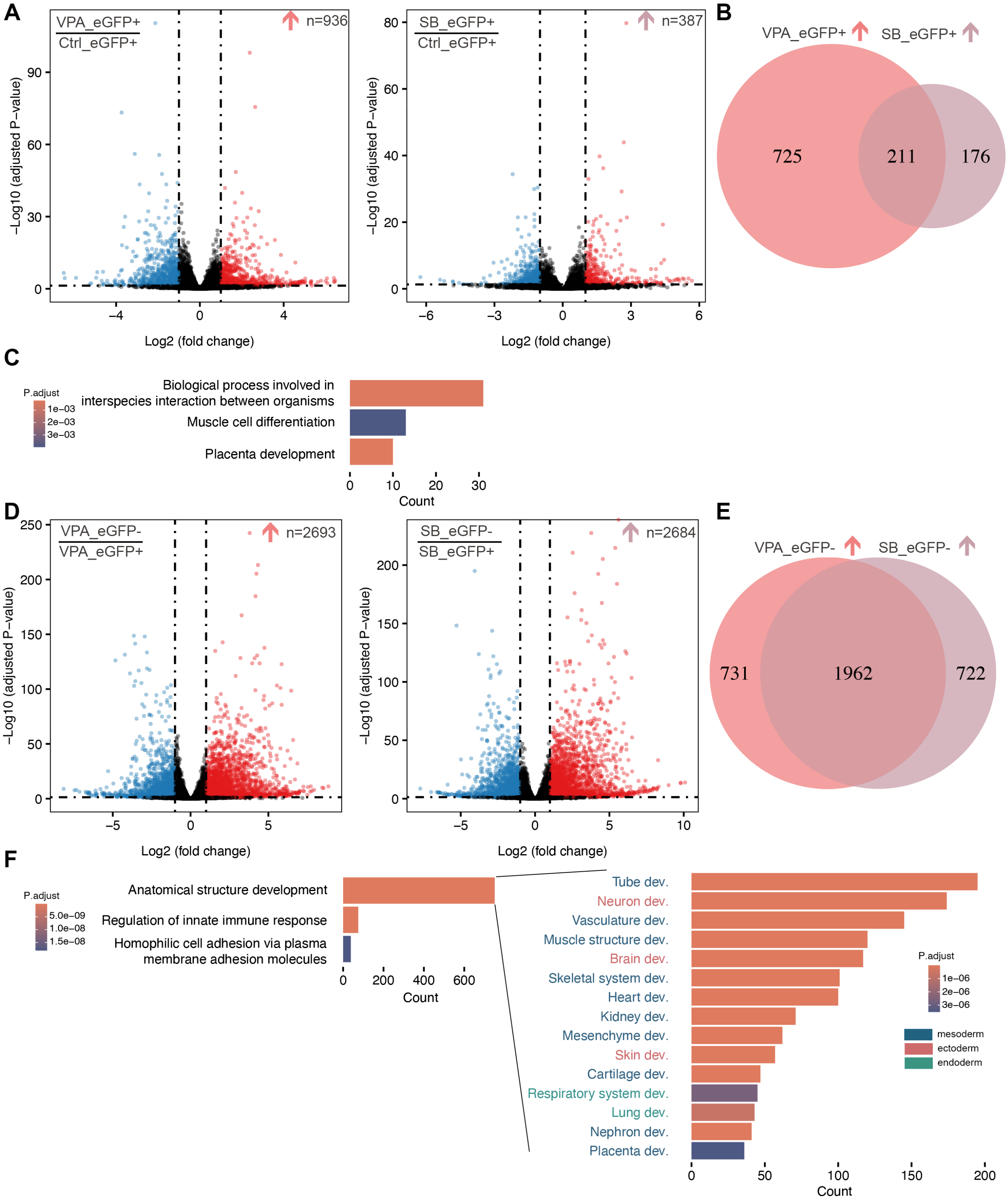


**Figure S16. Evidence of somatic lineage differentiation induced by HDACi treatment of differentiating PGCLC cultures.** A. Volcano plots of $\log_2$ (fold change) versus $-\log_{10}$ (adjusted P-value) for VPA_eGFP$^+$ (left) or SB_eGFP$^+$ (right) versus Ctrl_ eGFP$^+$ cells. B. Venn diagrams of the upregulated DEGs from A. C. Top enriched GO terms for the commonly upregulated genes in eGFP$^+$ cells in HDACi treatment groups. D. Volcano plots of Stella-eGFP$^-$ versus Stella-eGFP$^+$ cells treated with VPA (left) or SB (right). E. Venn diagram of the commonly upregulated DEGs from D. F. Left: Top enriched GO terms for the HDACi upregulated genes from E. Right: Count distribution of the “child” GO terms under “Anatomical structure development”.

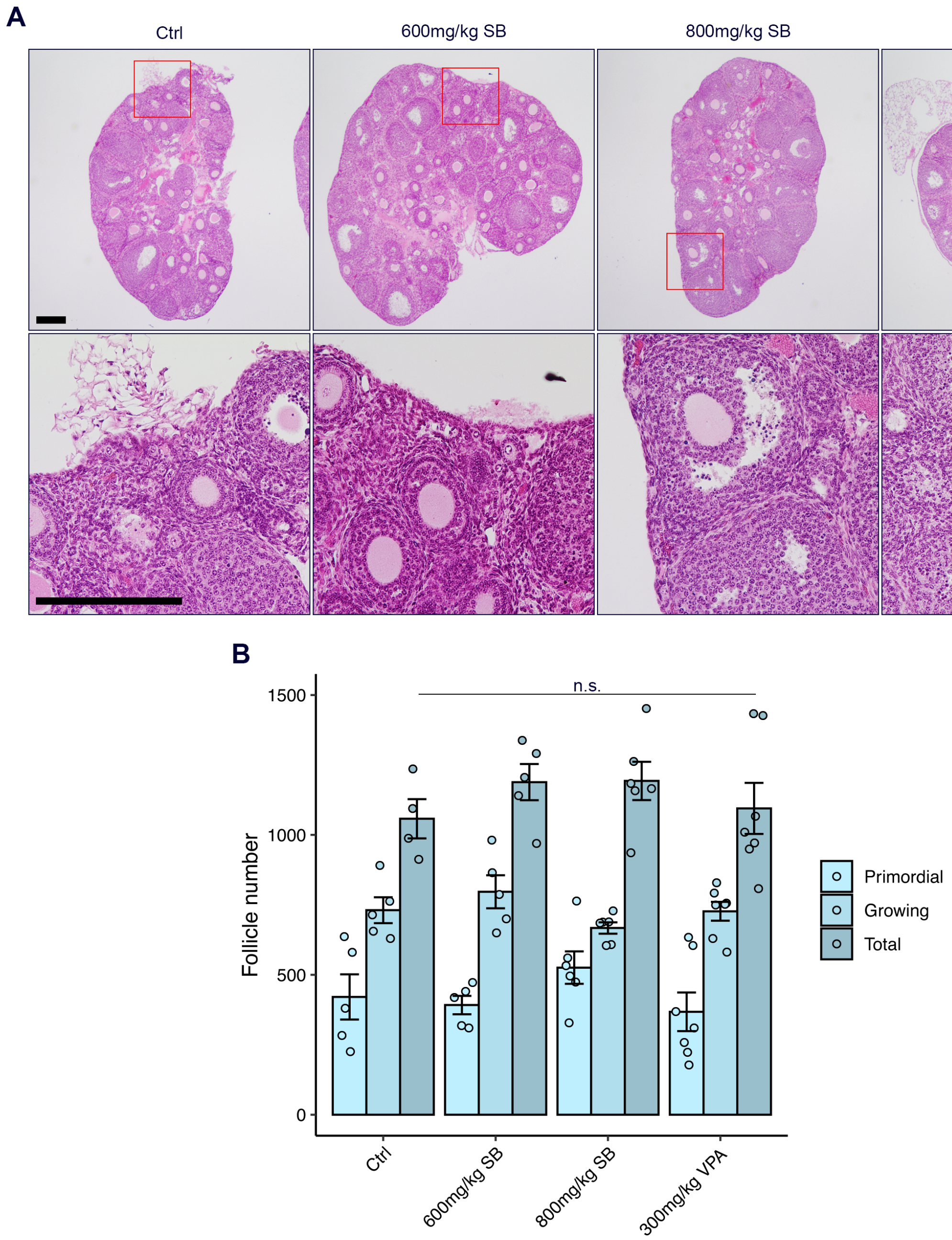


**Figure S17. Phenotypic analysis of female pups.** (A) Histological analyses of 1-month old ovaries. The boxed regions are magnifications of follicles showing in lower panel. Scale bars = 100 μm. (B) Follicle counts summed across every fifth serial section. Data are represented as the mean ± SEM and were analyzed using one-way ANOVA with Tukey's *post hoc* test. n.s. represents no significant difference.

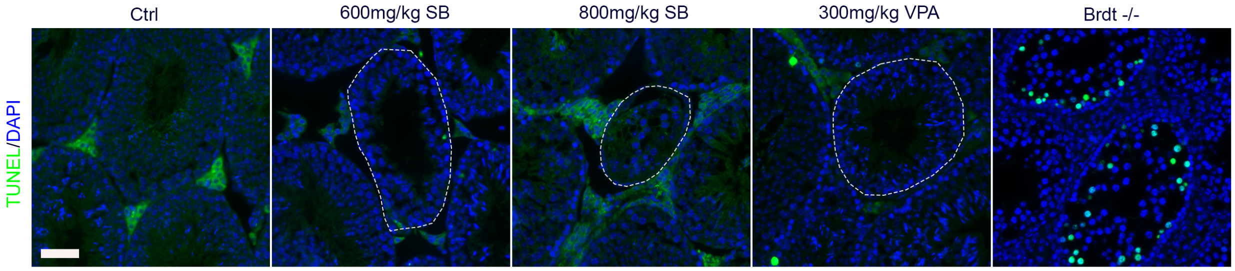


**Figure S18. Absence of apoptotic cells in atrophic seminiferous tubules.** Dashed lines denote regions with atrophic seminiferous tubules. Testicular section from *Brdt*$^{-/-}$ mouse model with a meiotic defect (Ding et al, 2023) was utilized as a positive control for TUNEL staining. Scale bar represents 50 µm.

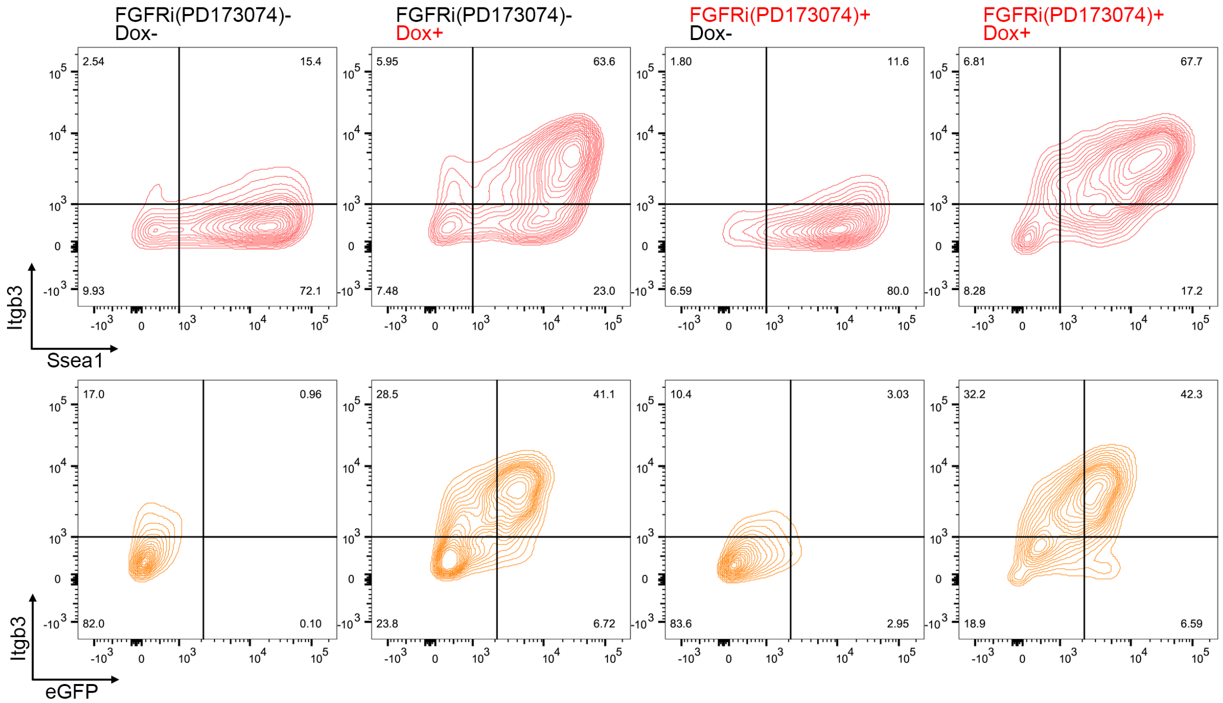


**Figure S19. Impact of FGFRi on differentiation of 4TF-induced PGCLCs from formative ESCs.** Day 6 cells from untreated 12-well plates were collected for FACS.

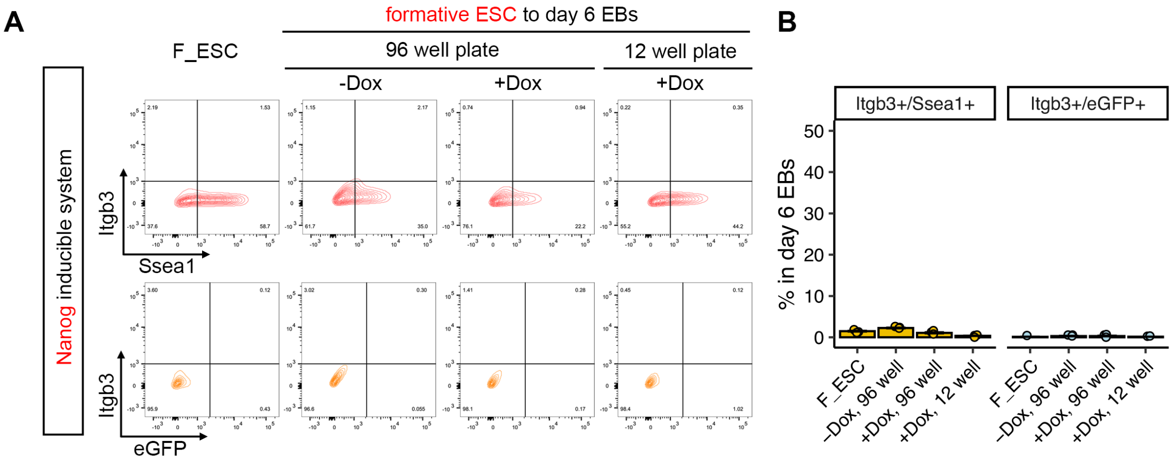


**Figure S20. Overexpression of Nanog in suspended formative ESCs.** A. Representative FACS pattern of Itgb3$^{+}$/Ssea1$^{+}$ and Itgb3$^{+}$/eGFP$^{+}$ cells in formative ESC and day 6 EBs from different plates and culture conditions. B. Quantification of PGCLC populations. Data in B are represented as the mean ± SEM.

| | Ck | 3TFs OE | Nanog OE | 4TFs OE |
|---|---|---|---|---|
| High efficiency | No | No | No | Yes |
| Scalability | No | N.A. | Yes | Yes |
| Long-term PGCLC culture | N.A. | N.A. | No | Yes |
| Direct diff. from formative ESC | Yes | N.A. | No | Yes |

**Figure S21. Comparation features of 4TF-inducible system with current available PGCLC differentiation systems.** N.A., not available.
