## Supplemental Text for "Scalable primordial germ cell-like-cell platform for functional genomics identifies epigenetic fertility modifiers"

### Supplementary Text

**HDACi increases expression of genes involved in somatic lineages**

We performed transcriptome analyses of eGFP$^+$ and eGFP$^-$ cells from day 6 control or HDACi-treated EBs (Fig. S15A). PCA revealed that day 6 VPA-treated Stella-eGFP$^-$ cells (VPA_eGFP$^-$) and SB_eGFP$^-$ cells clustered separately from the eGFP$^+$ cohorts (VPA_eGFP$^+$, SB_eGFP$^+$ and Ctrl_eGFP$^+$). However, amongst eGFP$^+$ cells, those treated with VPA clustered apart from the SB and control groups (Fig. S15B). k-Means analysis of top 2,000 DEGs and defined three clusters (C1, C2 and C3) (Fig. S15C). The genes in clusters C1 and C2, which were particularly elevated in eGFP$^-$ cells, were enriched with terms related to immune responses, cell migration and neural development, whereas cluster C3 containing predominantly eGFP$^+$ cells were enriched for genes involved in processes related to reproduction such as gamete generation and meiosis (Fig. S15C). GSVA revealed that eGFP$^-$ cells from both HDACi treated groups (VPA_eGFP$^-$ and SB_eGFP$^-$) displayed lower enrichment of Biological Processes related to reproduction and germline development (Fig. S15D). By contrast, eGFP$^+$ cells from control and treatment groups exhibited significant enrichment in all reproductive pathways, though the enrichment was greater in the untreated control group (Fig. S15D). Despite HDACi treatment, genes related to early PGC development were normally expressed in eGFP$^+$ cells. However, they were downregulated in eGFP$^-$ cells. For genes tied to meiosis and piRNA metabolic processing, their expression decreased under HDACi treatment, even in the eGFP$^+$ cells (Fig. S15E). These phenomena raise the possibility that failure to properly deacetylate (downregulate) non-germ cell regulatory elements may promote alternative lineage fates, a hypothesis tested by experiments described in the following section.

**HDACi treatment increases expression of genes involved in somatic lineages**

To further explore the identify and fate of cells exposed to HDACi, we focused on up-regulated DEGs in VPA_eGFP$^+$ and SB_eGFP$^+$ cells compared to Ctrl_eGFP$^+$ cells (Fig. S16A and B). Gene set enrichment analysis (FDR < 0.05) of 211 overlapping genes revealed clusters of upregulated GOBP driver terms in VPA_eGFP$^+$ and SB_eGFP$^+$ cells such as placental development, muscle cell differentiation, and innate immunity responses (Fig. S16C). The therapeutic effects of HDACi in compromising innate immunity and bacterial-viral entry may explain the observed GO terms such as innate immune response and defense response (1). The emergence of pathways related to placental and reproductive structure development and muscle and cytoskeleton development suggests possible incomplete suppression of factors involved in extraembryonic and embryonic mesoderm lineage differentiation.

Lacking precise knowledge of the mechanism governing mesoderm lineage activation with HDACi treatment, we hypothesized that $eGFP^+$ cells in treatment groups yield a transcriptome that is intermediate between germline and somatic fates. To test this, we applied gene set enriched analysis of up-regulated DEGs in HDACi-treated $eGFP^-$ cells compared to their $eGFP^+$ counterparts (Fig. S16D and E). The overlap genes enriched in GOBP driver terms such as anatomical structure development, regulation of innate immune response, and homophilic cell adhesion. The first category was enriched for GO terms associated with development of the placenta and all three germ layers including embryonic, cardiac, paraxial (muscle, skeleton, and cartilage), intermediate (kidney, nephron, and urogenital), and lateral (mesenchyme, blood vessel, and vasculature) mesoderm. Pathways associated with endoderm (lung, respiratory system) and ectoderm (skin and brain) development were also enriched (Fig. S16F). The results suggest that gene activation driven by histone hyperacetylation promotes overall somatic-lineage differentiation from epiblast cells.

**Discussion**

We opted to use a CRISPRi rather than a standard knockout-oriented CRISPR-Cas9 strategy, reasoning that complete ablation of some epigenetic genes would be generically disruptive to cell proliferation, and not representative of genetic or environmental perturbations. We identified 53 epigenetic candidates that compromised PGCLC formation, and those with the lowest beta scores (i.e., were most impaired in PGCLC differentiation) are normally upregulated in PGCLCs compared to naïve ESCs and EpiLCs, including *Rai1*, *Smarcd2*, *Kdm5d*, *Jak2*, *Ss18l1* and *Ncor2*. Knockdown of *Ncor2* was shown to inhibit the formation of $BLIMP1^+$ PGCLCs, suggesting an important role in mediating histone deacetylation in germ cell lineage specification (2). As a component of the nuclear receptor corepressor complex, *Ncor2* primarily contributes to gene repression through histone deacetylation. We confirmed *Ncor2*'s impact on

PGCLC development using siRNA to transiently suppress *Ncor2*, which led to the depletion of ITGB3$^{+}$/Stella-eGFP$^{+}$ PGCLCs. These results suggest that *Ncor2* plays a crucial role in suppression of transcriptional programs opposing the germline lineage fate. This gene suppression may be associated with the recruitment of Class I and Class IIa HDACs by the NCOR2 co-repressor complex. Previous work by Mochizuki et al. demonstrated that BLIMP1-HDAC3-mediated somatic gene repression is vital for PGC fate determination. Additionally, they showed that *Ncor2* knockdown caused BLIMP1$^{+}$ PGCLCs to have elevated expression of somatic lineage genes such as *Hoxb1*, *Hand1*, *Sox17*, and *Gata4* (2). We didn't identify individual Class I or Class IIa HDACs in our screen. These findings lead us to believe that *Ncor2* potentially inhibits somatic lineage development by interacting with Class I and Class IIa HDACs, and that there is redundancy amongst various HDACs.

The transcriptomes of Stella-eGFP$^{+}$ and Stella-eGFP$^{-}$ cells following HDACi treatment fell into distinct clusters by PCA, supporting Stella as a reliable marker for mouse PGC development. Indeed, downregulated genes in Stella-eGFP$^{-}$ cells (vs. Stella-eGFP$^{+}$) contained those related to reproduction, suggesting a suppression of cell fate. GSVA on reproductive pathways also indicated loss of germline fate even in the HDACi treated Stella-eGFP$^{+}$ PGCLCs, as indicated by relative depletion of PGC-associated transcripts such as *Nanos3* and *Dazl*. Interestingly, unperturbed Stella-eGFP$^{+}$ PGCLCs expressed several meiosis-related genes, and these meiotic genes were downregulated in the HDACi treated cells. Even though there was no indication that the differentiated cells were differentiating into meiocytes, the PGCLCs may be partially inducing a meiotic transcriptional program that HDACi treatment disrupts. Additionally, HDACi-treated Stella-eGFP$^{+}$ PGCLCs exhibited upregulation of pathways associated with mesoderm development, and there was significant upregulation of GO terms related to somatic lineage development in Stella-eGFP$^{-}$ cells. In mice, the induction *Prdm1* and *Prdm14* is crucial for driving PGC fate by suppressing mesoderm lineage genes (3).

The coordinated regulation of BMP and WNT/β-Catenin signaling is pivotal for the specification of mouse PGCs from the proximal epiblast (4). However, signaling through WNT/β-Catenin alone is insufficient germline commitment. While the activation of

WNT/β-Catenin does enhance the expression of T, it simultaneously triggers numerous mesoderm transcriptional regulators such as *Sp5*, *Hoxa1*, *Hoxb1*, *Nkx1-2*, *Msx2*, and *Rreb1* (5). The exposure to BMP4 of cells in the proximal epiblast suppresses these mesoderm factors, creating an optimal environment for T to activate *Prdm1* and *Prdm14*, initiating the emergence of PGCs. In HDACi-treated Stella-eGFP$^{-}$ cells, we noted upregulation of these mesoderm regulators, but downregulation of *Ifitm3,* which is activated in PGCs by WNT3A and BMP4 signaling (6). This suggests suppression of germline fate in the presence of elevated signaling from other lineage differentiation factors. These findings support the idea that exposure to HDACi can negatively impact germline development.

Besides *Ncor2*, our screen identified several other candidate epigenetic regulators of PGC development. These include BLIMP1, which suppresses somatic gene expression (7), and TLE1, which interacts with the BLIMP1 repression domain to function as a co-repressor complex in B cells (8). Loss of *Tle1* is linked with elevated acetylation levels and gene activation (9), suggesting a role analogous to *Ncor2* in suppressing somatic gene expression in the germline. Other hits included Janus kinase 2 (JAK2), the absence of which causes a loss of gonadal PGCs in chicken (10), and BMI1, an element of the Polycomb repressive complex 1 (PRC1) that is vital for the preservation of spermatogonial stem cells (11). However, its significance in prenatal germ cells has not been elucidated. Focused studies of these and other genes identified in the screen will be needed to validate and understand their roles.
